# Toxicity potentials of selected insecticidal plant oils against *Anopheles gambiae* (Diptera: Culicidae) under laboratory conditions

**DOI:** 10.1101/2022.08.14.503910

**Authors:** Olukayode James Adelaja, Adedayo Olatubosun Oduola, Adeolu Taiwo Ande, Oyindamola Olajumoke Abiodun, Abisayo Ruth Adelaja

**Affiliations:** Department of Zoology, University of Ilorin, Ilorin, Nigeria; Department of Chemistry, University of Ilorin, Ilorin, Nigeria; Department of Pharmacology and Therapeutics, University of Ibadan. Ibadan, Nigeria

**Keywords:** *Anopheles gambiae*, Plant oils, Toxicity, Mortality

## Abstract

Despite increasing reports and concerns about resistance development to public-health insecticides in malaria-vectors, significant steps have been put into the quest for novel strategies to disrupt the disease transmission cycle by targeting insect-vectors hence sustaining vector management. This study evaluates the toxicity potential of oils of insecticidal plants shortlisted in an ethnobotanical survey on the larvae and adult stages of *Anopheles gambiae.* Oils from leaves of *Hyptis suaveolens, Ocimum gratissimum, Nicotiana tabacum, Ageratum conyzoides* and fruit-peel of *Citrus sinensis* were extracted by steam-distillation using a Clevenger apparatus. Larvae and female adults of deltamethrin susceptible *Anopheles gambiae* were gotten from an already established colony in the Entomological Research Laboratory, University of Ilorin. Twenty-five third instar stage larvae were used for larvicidal assays while twenty 2-5 days old adults were used for the adulticidal assays in five replicates. After which, coupled gas chromatography-mass spectrometry analysis (GC-MS) was performed to determine the major chemical-constituents of plant oils. *A. gambiae* exposed to *H. suaveolens* and *C. sinensis* demonstrated significantly higher larval toxicity (94.7-100%) after 24 hours. At 48 hours, the mortality induced by the oils of the four plants peaked at 100%. *N. tabacum* (0.50 mg/ml) induced the highest percentage adult mortality (100%) on *A. gambiae* which compared favourably with the positive control Deltamethrin (0.05%). The lowest KdT_50_ was observed with 0.25 mg/ml of *N. tabacum* (20.3 minutes) while the lowest KdT_95_ was observed with 0.10mg/ml of *A. conyzoides* (35.97 mins) against adult *A. gambiae.* D-limonene is the key chemical-constituent in oils from *C. sinensis* and *A. conyzoides.* The significant larval and adult mortality rates, lower lethal concentration and knockdown times demonstrated by the evaluated plant oils showed promising outcomes that can be further developed for vector control management.

**Graphical Abstract:** 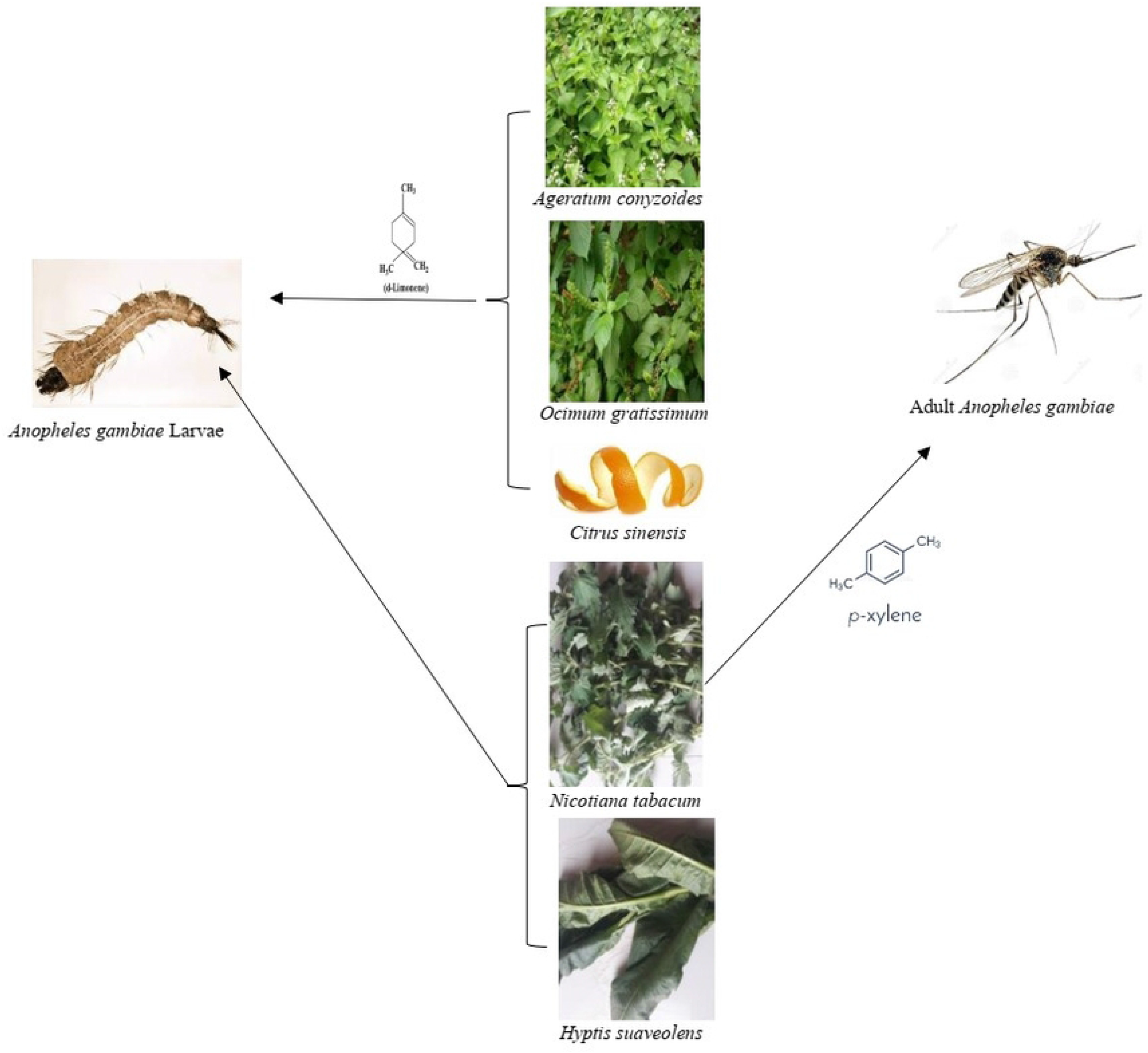

## Introduction

Mosquito-borne disease burdens accounts for seventeen percent of all infectious diseases and seven hundred thousand deaths yearly (WHO 2019a). Malaria, whose mosquito vector is *Anopheles* spp., frank the highest with approximately 229 million cases in 2021 resulting in 409,000 deaths globally, 95% of which occurred in Africa and 23% in Nigeria. (WHO 2021)

In Nigeria, *Anopheles gambiae* s.l. (Giles); is a complex with numerous sibling species including *A. gambiae, A. arabiensis, A. coluzzii, A. quadriannulatus, A. amharicus, A. bwambae, A. merus* and *A. melas* have been documented as malaria vectors. The first three sibling species have however been incriminated as responsible for the high malaria transmission, morbidity and mortality statistics in Nigeria (Okorie et al. 2011; Olayemi et al. 2011; Oduola et al. 2016; Obi et al. 2017).

The prime choice mosquito control tactic in Africa remains the selective application of residual synthetic insecticides through either Indoor Residual Spraying (IRS) or Long-Lasting Insecticide Nets (LLIN) use (WHO 2019a). To this end, Pyrethroids are members of the prequalified classes of active ingredient of World Health Organisation (WHO) vector control product (Omotayo et al. 2021; WHO 2021; Ande et al. 2022). However, insecticide resistance development in mosquitoes poses a serious threat to sustainable synthetic insecticide-based vector control programmes in many African countries (Youmsi et al. 2017; Oduola et al. 2019) and necessitates the search for natural, sustainable and eco-friendly alternatives.

Cultural practices over the years coupled with research outputs have shown that plants are rich sources of active constituents that have been used successfully and persistently against mosquitoes (Youmsi et al. 2017; Ogban et al. 2020; Adelaja et al. 2021). In many rural communities in Africa, plants are being used efficiently for the management of insects of concern including mosquitoes (Nzelibe and Chintem 2015; Karunamoorthi et al. 2009) and they have been reported to be readily accessible, available, affordable and biodegradable (Kamaraj et al. 2010). Plants, therefore, constitute a potential source of natural and sustainable agents that will stand the test of time as a mosquito control agent.

In Nigeria, numerous plants and plant parts have been reportedly used in the management of larvae or adult mosquitoes across various cultures and over the years (Ivoke et al. 2009; Mgbemena 2010; Okigbo et al. 2010; Ileke et al. 2015; Musa et al. 2015; Nzelibe and Chintem 2015; Ayange-Kaa et al. 2015; Owoeye et al. 2016; Adelaja et al. 2021). The potentials and modes of actions of these plants have been scientifically validated through research reports (Ogban et al. 2020; Adelaja et al. 2021). Crude extracts of some of such plants reported include *Ageratum conyzoides* (L.), *Hyptis suaveolens* (L.), *Ocimum gratissimum* (L.) and *Nicotiana tabacum* (L.) (Ivoke et al. 2009; Mgbemena 2010; Okigbo et al. 2010; Ileke et al. 2015; Musa et al. 2015; Nzelibe and Chintem 2015; Ayange-Kaa et al. 2015; Owoeye et al. 2016; Adelaja et al. 2021). There is, however dearth of knowledge in respect of the full compliments of the chemical constituents and the toxicity potentials of the oils obtained from each of these botanicals and *Citrus sinensis* (L.) against *A. gambiae.* This study, therefore, provides valuable baseline information on the major chemical constituents and the toxicity potentials of each of the above-mentioned plant oils on *A. gambiae.*

## Materials and Methods

### Plant Materials, Oil Extraction and Serial Dilution Procedures

The leaves of *Hyptis suaveolens* (Lamiaceae)*, Ocimum gratissimum* (Lamiaceae)*, Nicotiana tabacum* (Solanaceae)*, Ageratum conyzoides* (Asteraceae) and fruit peel of *Citrus sinensis* (Rutaceae) previously identified as most frequently used insecticidal plants from an ethnobotanical survey carried out among locals from different villages in North-central Nigeria (Adelaja et al. 2021) were sourced individually from fresh growing plants around Ilorin. Each collection was identified at the herbarium of the Plant Biology Department by accessory detail and air dried under shade for one week before hydro-distillation. The respective plant oils obtained from each pulverized plant sample were isolated by steam-distillation using a Clevenger apparatus. Each isolated oil was dried over anhydrous sodium sulphate and stored in amber-coloured vials at 4°C until required for subsequent assays. Each of the five plant oils to be evaluated were serially diluted using technical grade acetone into eight concentrations, viz: 5% (vol:vol) – 0.05mg/ml, 10% (vol:vol) – 0.10 mg/ml, 15% (vol:vol) – 0.15 mg/ml, 20% (vol:vol) – 0.20mg/ml, 25% (vol:vol) – 0.25 mg/ml, 30% (vol:vol) – 0.30 mg/ml, 40% (vol:vol) – 0.40 mg/ml and 50% (vol:vol) – 0.50 mg/ml. These concentrations were determined based on previously unpublished data.

### Chemical Constituents Assay

Coupled gas chromatography-mass spectrometry analysis (GC-MS) of the extracted plant oils was performed on a Hewlett Packard 5890 II gas chromatograph, interfaced to a single quadrupole mass selective detector (Model 5972). The column was an HP-5 MS capillary column (30 × 0.25 mm, film thickness 0.25 mm). Helium was the carrier gas, set at a flow rate of 0.6 ml/min. Injector and MS transfer line temperatures were set at 220 and 250 °C, respectively. The oven programme temperature was held at 35 °C for 5 mins and then 4 °C/min to 150 °C for 2 mins and then finally 20 °C/min to 250 °C for 5 mins. Diluted samples (10:100 in CH_2_Cl_2_, v/v) of 1 μL were injected manually and in a split mode (1:100 split ratio). The identification of the components was accomplished by comparison of their relative retention indices as well as comparison of mass spectra with those of standards, those found in the literature and those supplemented by the National Institute of Standards and Technology (NIST) provided by Hewlett Packard with the GC/ MS control and data processing software (Joshi 2017).

### Mosquito Collection and Rearing

The larvae and female adults of deltamethrin susceptible *Anopheles gambiae* mosquitoes were gotten from an already established colony at the insectary of the Entomological Research Laboratory, University of Ilorin which was maintained at 27±20°C and 85±5% RH. Mosquito larvae were fed a diet of non-fatty biscuits mixed with yeast at a ratio of 3:1 while emerged adults were confined in adult holding cages (50 x 50 x 50cm) and provided with a 10% sugar solution. Adult mosquitoes were sorted into males and females using morphological keys by Gillies and DeMeillon (1987).

### Larvicidal Bioassay

The larvicidal activity of each of the plant oils at each of the serial dilution concentrations was evaluated according to the protocol of the World Health Organisation (WHO 2005) with slight modifications. Twenty-five (25) third instar larvae of *A. gambiae* mosquitoes were introduced into test cups containing 100ml of distilled water and allowed to acclimatise for 1hour. After which 2ml of the various plant oil concentrations described earlier were introduced. Distilled water with and without acetone was used as negative and positive controls respectively. The number of dead larvae was recorded after 24 and 48 hours of exposure and the percentage mean mortality was calculated. The larvae were considered dead if they did not move when prodded with a needle in the siphon or cervical region. Each treatment was tested in five replicates. If the negative control mortality was between 5% and 20%, the mortalities of treated groups were corrected according to Abbott’s formula (Abbott 1925).

### Adulticidal Bioassay

The adulticidal activity of the plant oils was evaluated against female adults *A. gambiae* mosquitoes following the World Health Organisation standard method (WHO 2013) with slight modifications. Two and a half millilitres (2.5 ml) of oil dilution concentration to be tested was impregnated on 12 × 15 cm Whatman paper (Dua et al. 2008). Deltamethrin (0.05%) was used as the positive control while acetone was used as the negative control. The impregnated papers were air-dried for 5 min and then inserted to line the interior of the exposure tube in the WHO test kit. Twenty (20), 2-5-day-old, blood-starved adult female mosquitoes were introduced into the holding tube and held for 1h to acclimatize. The mosquitoes were transferred gently into the exposure tube and knocked down was monitored and determined for 60 minutes and percentage knockdown was calculated afterwards. After 1 hour in the exposure tube, all mosquitoes were gently transferred back to the holding tube for recovery. A small ball of cotton wool soaked with 10% glucose solution was placed on the mesh screen for recovering mosquitoes to feed on. The number of dead mosquitoes was recorded at the end of the 24 hours recovery period, and the percentage of mortality was calculated. The bioassay was replicated five times. If the negative control mortality was between 5% and 20%, the mortalities of treated groups were corrected according to Abbott’s formula (Abbott 1925).

### Data Analysis

Percentage knockdown and mortality were calculated using the formulae below:

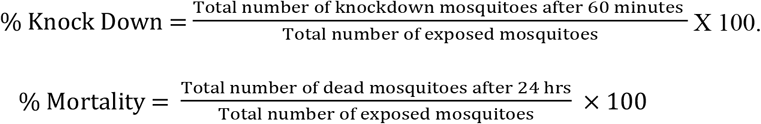

Analysis of Variance (ANOVA) and the Tukey post hoc test was used to compare the knockdown and mortality rates in test treatments and control using GraphPad Prism 8 software. The knockdown time for 50% (KdT_50_) and 95% (KdT_95_) of the adult *A. gambiae* population, as well as, the lethal concentration that can cause mortality 50% (LC_50_) and 90% (LC_90_) for the concentrations tested (0.05, 0.10, 0.15, 0.20, 0.25, 0.30, 0.40, 0.50 mg/ml) of the larvae of *An. gambiae* population were determined using probit analysis (Finney 1971) with the aid of GraphPad Prism 8 software.

## Results

### Percentage composition of major chemical constituents of the extracted plant oils

The major chemical constituents in the extracted plant oil from *Hyptis suaveolens, Citrus sinensis, Ocimum gratissimum, Nicotiana tabacum* and *Ageratum conyzoides* are shown in Table 1. It was discovered that D-Limonene was a common chemical constituent found in three of the five plant oils evaluated namely; *C. sinensis* (92.5%), *A. conyzoides* (63.96%) and *N. tabacum* (2.62%) and it was a major chemical constituent in the first two. Meanwhile, 3-Octen-1-ol was a chemical constituent found in *H. suaveolens* (1.09%) and *C. sinensis* (0.97%). Precocene I (95.63%) was the major constituent found in *O. gratissimum* and Caryophyllene (0.92%) was also found there while P-Xylene (12.37%) and Farnesol (3.08%) were the major constituents found in *N. tabacum*.

**Table 1.** Percentage composition of major chemical constituents of selected insecticidal plant oils.

| s/n | Component | *Hyptis*<br>*suaveolens* | *Citrus*<br>*sinensis* | *Ocimum*<br>*gratissimum* | *Nicotiana*<br>*tabacum* | *Ageratum*<br>*conyzoides* |
| --- | --- | --- | --- | --- | --- | --- |
| 1. | D-Limonene |  | 92.5 | 2.62 |  | 63.96 |
| 2. | 3-Octen-1-ol, (Z) | 1.09 | 0.97 |  |  |  |
| 3. | Precocene I |  |  | 95.63 |  |  |
| 4. | Caryophyllene |  |  | 0.92 |  |  |
| 5. | 1-Octyn-3-ol | 9.81 |  |  |  |  |
| 6. | P-Xylene |  |  |  | 12.37 |  |
| 7. | Farnesol |  |  | 0.05 | 3.08 |  |
| 8. | Preg-4-en-3-one |  |  | 0.05 |  | 0.58 |
| 9. | Paromomycin | 1.20 |  |  |  |  |

### Larvicidal Activity

Table 2 shows the mean larval mortality rates of *A*. *gambiae* exposed to various concentrations of each of the extracted plant oils and their LC_50_. and LC_90_ values. All plant oils were distinctly potent as toxicants against the third instar larvae with statistically different mortality rates (P<0.05) even at the lowest plant oil application concentration of 0.05mg/ml. It was also observed that toxicity intensity was concentration dependent meaning as concentration increased, mortality rates increased.

**Table 2.**
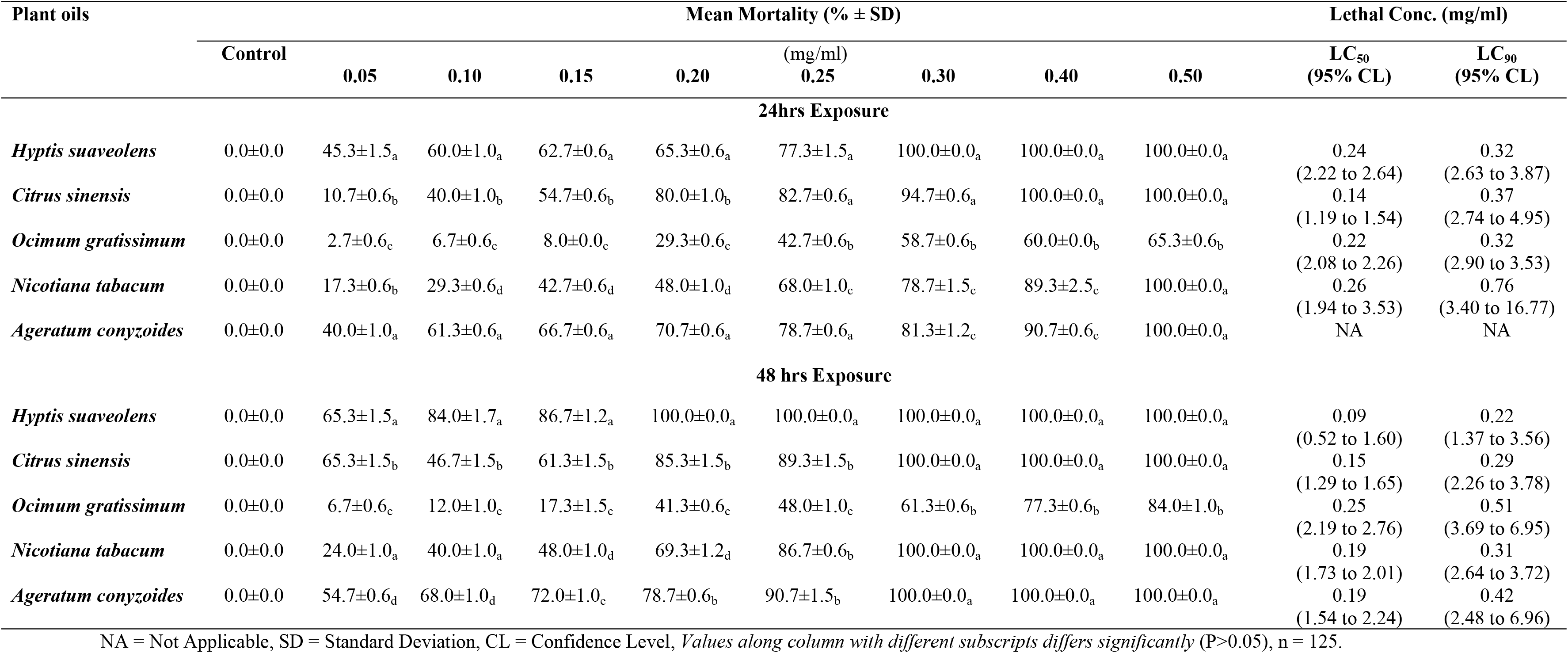
Mean *An. gambiae* larval mortality (induced by) recorded at various concentrations of some extracted plant oils and their LC_50_. and LC_90_ values.

After 24-hour exposure period, there was no significant difference (P>0.05) in the mean larval mortalities of the *A. gambiae* mosquitoes exposed to *H. suaveolens* (100%, 100%, 100%) and *C. sinensis* (94.7%, 100%, 100%), and also between *N. tabacum* (78.7%, 89.3%, 100%) and *A. conyzoides* (81.3%, 90.7%, 100%) at 0.30, 0.40 and 0.50 mg/ml respectively. However, the mean larval mortalities in *A. gambiae* mosquitoes exposed to these two groups of plant oils were significantly higher (P<0.05) when compared to mortalities (58.7%, 60.0% and 65.3%) observed in *A. gambiae* mosquitoes exposed to *O. gratissimum* at the three concentrations. (Table 2).

After 48-hour exposure period, mean larval mortality was 100% in *A. gambiae* exposed to 0.20 mg/ml of *H. suaveolens,* 0.30 mg/ml of *C. sinensis, N. tabacum* and *A. conyzoides.* The highest mortality (84.0%) in *A. gambiae* mosquitoes exposed to *O. gratissimum* occurred at 0.50 mg/ml. The percentage mortality observed in *A. gambiae* mosquitoes exposed to *H. suaveolens* at 0.20 mg/ml was significantly higher (P<0.05) when compared to mortalities observed in *A. gambiae* mosquitoes exposed to *C. sinensis, N. tabacum* and *A. conyzoides* (85.3%, 69.3%, 78.7% respectively) at same concentration. The mean larval mortalities of *A. gambiae* mosquitoes exposed to *O. gratissimum* was significantly lower (P<0.05) to that of other plant oils at all concentrations (Table 2).

The LC_50_ measurements indicated that *C. sinensis* (0.14 mg/ml) was more toxic at 24 hours exposure compared to *O. gratissimum* (0.22 mg/ml), *H. suaveolens* (0.24 mg/ml) and *N. tabacum* (0.26 mg/ml). However, at both 24- and 48-hours exposure, *H. suaveolens* showed lower LC_90_ (0.32 mg/ml, 0.22 mg/ml) larval toxicity when compared with oils of *C. sinensis* (0.37 mg/ml, 0.29 mg/ml), *N. tabacum* (0.76 mg/ml, 0.31mg/ml), *A. conyzoides* (NA, 0.42 mg/ml) and *O. gratissimum* (0.32 mg/ml, 0.51 mg/ml) (Table 2).

### Adulticidal Activity

Figure 1 shows the adulticidal activity of various concentrations of the extracted plant oils against *A*. *gambiae* after 24 hrs exposure and Table 3 shows percentage knockdown after one hour and their LC_50_. and LC_90_ values. All plant oils were distinctly potent as toxicants against the female adult *A*. *gambiae* with statistically different mortality rates (P<0.05) even at the lowest plant oil application concentration of 0.05 mg/ml. It was also observed that toxicity intensity was concentration dependent meaning as concentration increased, mortality rates increased also (Figure 1).

**Fig. 1.**
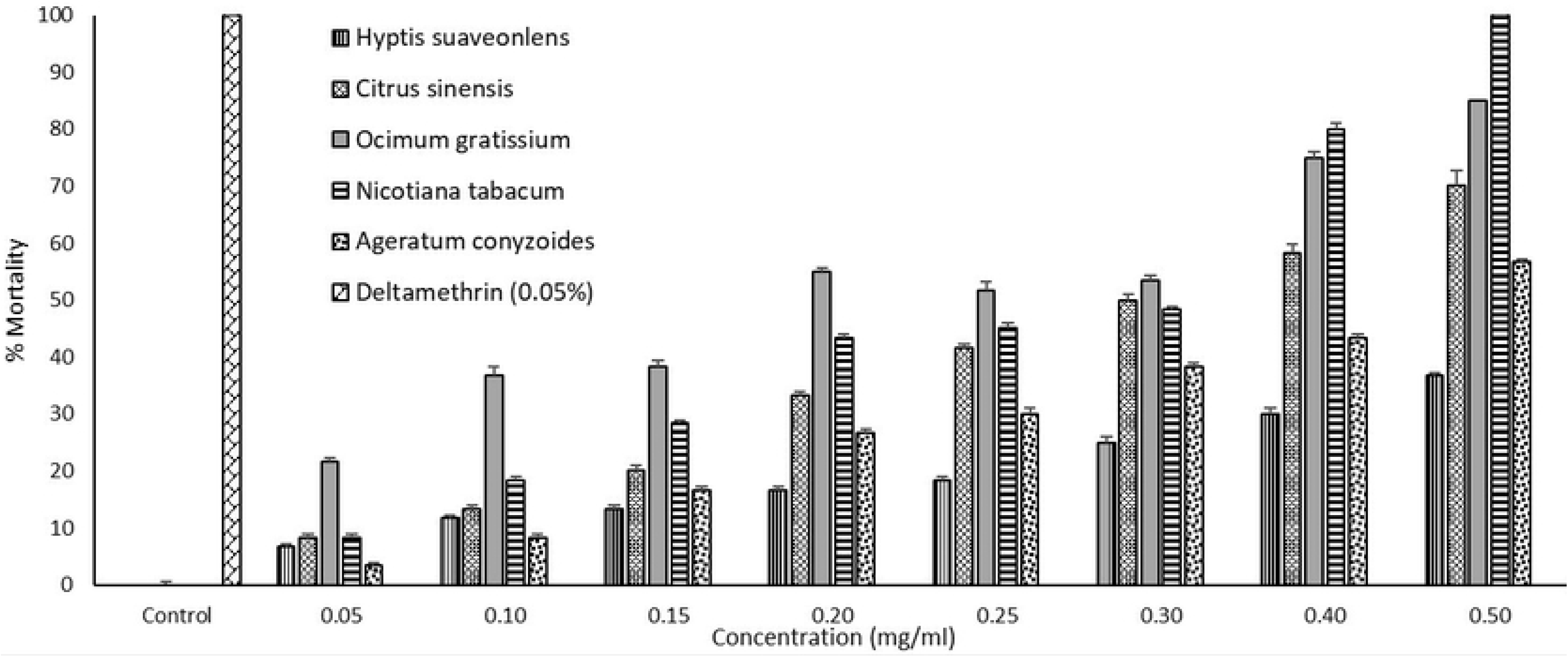
Adulticidal activity (Dose-response assay) of selected plant oils on *Anopheles gambiae* after 24 hrs post exposure.

*N. tabacum* at 0.50 mg/ml induced the highest percentage mortality (100%) on *A. gambiae* which compared favourably with Deltamethrin (0.05%), a positive experimental control insecticide. Also, exposure of *A. gambiae* to *O. gratissimum* at O. 40 and 0.50 mg/ml and *N. tabacum* at 0.40 mg/ml) resulted in a minimum of 75%, 85% and 80% mortalities, respectively. The lowest percentage of mortality was recorded in 0.05 mg/ml of *A. conyzoides* (3.3%) (Figure 1).

The lowest KdT_50_ was recorded with *N. tabacum* at 0.25 mg/ml (20.3 mins) followed by *O. gratissimum* at 0.30 mg/ml (22.93 mins), while the highest KdT_50_ was recorded with *C. sinensis* at 0.05 mg/ml (75.56 mins) followed by *H. suaveolens* at 0.30 mg/ml (73.25 mins). Meanwhile, the lowest KdT_95_ was recorded with *A. conyzoides* at 0.10 mg/ml (35.97 mins) and 0.05 mg/ml (45.27 mins) followed by *O. gratissimum* at 0.30 mg/ml (48.11 mins) while the highest was recorded with *N. tabacum* at 0.20 mg/ml (313.1 mins) followed by *C. sinensis* at 0.05 mg/ml (288.9 mins) and 0.30 mg/ml (267.5 mins) (Table 3).

**Table 3.** Knock-down Effect and Knockdown times ratings of selected plant oils against *Anopheles gambiae*.

| *Plant oils* | **Conc.**<br>**(mg/ml)** | **% Kd after**<br>**1hr** | **KdT<sub>50</sub> (95%CL)** | **KdT<sub>95</sub> (95%CL)** |
| --- | --- | --- | --- | --- |
| *Hyptis suaveolens* | 0.05 | 6.67 | 31.05 (22.80 to 42.29) | 53.88 (19.30 to 150.4) |
|  | 0.10 | 11.67 | 49.86 (24.19 to 102.8) | 102.3 (19.65 to 532.4) |
|  | 0.15 | 13.33 | NA | NA |
|  | 0.20 | 16.67 | NA | NA |
|  | 0.25 | 18.33 | NA | NA |
|  | 0.30 | 25.00 | 73.25 (NA) | NA |
|  | 0.40 | 30.00 | NA | NA |
|  | 0.50 | 36.67 | NA | NA |
| *Citrus sinensis* | 0.05 | 8.33 | 75.56 (0.06 to 93831) | 288.9 (NA) |
|  | 0.10 | 13.33 | NA | NA |
|  | 0.15 | 20.00 | NA | NA |
|  | 0.20 | 33.33 | NA | NA |
|  | 0.25 | 41.67 | NA | NA |
|  | 0.30 | 50.00 | 48.19 (0.9433 to 2462) | 267.5 (NA) |
|  | 0.40 | 58.33 | 29.06 (25.49 to 33.13) | 48.23 (32.33 to 71.93) |
|  | 0.50 | 70.00 | 36.21 (20.93 to 62.63) | 84.95 (14.79 to 488.1) |
| *Ocimum gratissimum* | 0.05 | 21.67 | NA | NA |
|  | 0.10 | 36.67 | NA | NA |
|  | 0.15 | 38.33 | 37.08 (23.97 to 37.22) | 121 (15.01 to 975.0) |
|  | 0.20 | 55.00 | 40.1 (18.90 to 85.05) | 144.5 (16.65 to 1254) |
|  | 0.25 | 51.67 | 34.19 (20.35 to 57.41) | 152.5 (22.11 to 1053) |
|  | 0.30 | 53.33 | 22.93 (18.84 to 27.91) | 48.11 (25.77 to 89.83) |
|  | 0.40 | 75.00 | NA | NA |
|  | 0.50 | 85.00 | NA | NA |
| *Nicotiana tabacum* | 0.05 | 8.33 | 30.90 (20.51 to 46.56) | 69.21 (16.84 to 284.5) |
|  | 0.10 | 18.33 | NA | NA |
|  | 0.15 | 28.33 | 29.87 (19.01 to 72.31) | 77.19 (34.32 to 173.6) |
|  | 0.20 | 43.33 | 60.4 (4.634 to 787.4) | 313.1 (1.547 to 63336) |
|  | 0.25 | 45.00 | 20.3 (8.496 to 48.49) | 98.22 (4.662 to 2069) |
|  | 0.30 | 48.33 | 32.96 (25.57 to 42.47) | 78.26 (32.88 to 186.3) |
|  | 0.40 | 80.00 | NA | NA |
|  | 0.50 | 100.00 | 27.89 (23.15 to 33.60) | 69.74 (35.07 to 138.7) |
| *Ageratum conyzoides* | 0.05 | 3.33 | 44.68 (NA) | 45.27 (NA) |
|  | 0.10 | 8.33 | 26.29 (19.79 to 34.93) | 35.97 (23.43 to 55.22) |
|  | 0.15 | 16.67 | NA | NA |
|  | 0.20 | 26.67 | NA | NA |
|  | 0.25 | 30.00 | 64.02 (13.25 to 309.4) | 129.1 (9.150 to 1821) |
|  | 0.30 | 38.33 | NA | NA |
|  | 0.40 | 43.33 | 30.25 (25.76 to 35.52) | 69.59 (39.76 to 121.8) |
|  | 0.50 | 56.67 | NA | NA |

Conc. =Concentration, % Kd = Percentage KnockDown, NA = Not Applicable, KdT = Knock Down Time, CL = Confidence limit, n = 100

## Discussion

Mosquito larvicidal efficacy of plant oils and extracts varies according to plant species, the part of the plant, the geographical location where the plant is grown and application methods (Ghosh and Chowdhury 2012; Adelaja et al. 2021). The current study investigated the toxicity potentials of the following shortlisted plants: *H. suaveolens, C. sinensis, O. gratissimum, N. tabacum* and *A. conyzoides* against a major malaria vector, *A. gambiae*.

The 100% *A. gambiae* larvae mortally recorded with 0.30 mg/ml concentration of *H. suaveolens,* 0.40 mg/ml concentration of *C. sinensis* and 0.50 mg/ml concentrations of *N. tabacum* and *A. conyzoides* after 24 hours as well as 0.20 mg/ml concentration of *H. suaveolens* and 0.30 mg/ml concentrations of *C. sinensis, N. tabacum* and *A. conyzoides* after 48 hours compared favourably with Kavendan et al. (2014) and Bobbo et al. (2016) where *H. suaveolens* caused high mortalities against *A. gambiae* mosquito larvae after 24 hours post-exposure. Similarly, Musa et al. (2015) reported that *A. conyzoides* evoked 92.0% and 87.5% mortality respectively against *A. gambiae* after 24hrs of exposure while Ileke et al. (2015) reported that *N. tabacum* elicited high mortality to the larvae of *A. gambiae*. Hence, oils from *H. suaveolens, C. sinensis*, *N. tabacum* and *A. conyzoides* could be considered and adopted as natural larvicides for the management of *Anopheles* mosquitoes in malarious areas where breeding sites can easily be identified and accessed. However, it was observed that all the concentrations of *O. gratissimum* oils tested elicited lower larvicidal activities against *A. gambiae* larvae and may therefore not be a promising larvicidal agent. Meanwhile, the lowest LC_50_ and LC_90_ (after 24- and 48-hours exposure) against *A. gambiae* larvae were recorded in oil from *C. sinensis* and *H. suaveolens*. Oils from *C. sinensis* and *H. suaveolens* displayed the highest mortality at the lowest concentrations echoing their potential as good larvicides that should be considered for the management of *Anopheles* mosquitoes locally.

In this study, *N. tabacum* at 0.50 mg/ml recorded a hundred percent mortality within a one-hour test period on adult *A. gambiae* comparing favourably with the Deltamethrin positive control. Similarly, Owoeye et al. (2016) and Ileke et al. (2015) have reported that the toxicity of *N. tabacum* against adult *A. gambiae* within a one-hour of test period. The lowest KdT_50_ against *A. gambiae* was recorded in 0.25 mg/ml of *N. tabacum* while the lowest KdT_95_ was recorded in 0.30 mg/ml of *O. gratissimum.* Therefore, the active ingredients in oil from *N. tabacum* can further be investigated and developed as alternatives to the use of pyrethroids for vector control purposes. Also, the high mortalities recorded in *A. gambiae* mosquitoes exposed to 0.50 mg/ml of *O. gratissimum* demonstrated its potential as alternative for *Anopheles* mosquito control in places where they naturally grow. This suggests that *N. tabacum* and *O. gratissimum* plant oils have the potential to be used as rapid knock down agent to be incorporated into aerosols used indoors for personal protection against adult endophagic and arthropogenic malaria vectors.

Further studies on the insecticidal plants being evaluated for their toxicity potential have identified some compounds with bioactivity. Some similar exercises have led to the isolation and characterization of the active constituents (Okwute 2012). In this study, D-Limonene was identified by GC-MS analysis as a chemical constituent in oils from *C. sinensis, A. conyzoides* and *O. gratissimum*. Ahmad et al. (2006) and Espina et al. (2011) also reported D-Limonene as a key chemical constituent in oils from *C. sinensis, A. conyzoides* and *O. gratissimum* even though Usman et al. (2013) identified it as the least abundant chemical constituent in oils from *A. conyzoides*. This however contrast with the outcome of this study indicating that D-Limonene was relatively abundant in *A. conyzoides*. However, it was the least abundant in *O. gratissimum.* The isolated forms of D-Limonene have been reported to have insecticidal potential against stored grain insects (Lee et al., 2003), cockroaches (Karr and Coats 1988) and termites (Almeida et al. 2015). D-Limonene has also been reported to act as a contact poison against some insects e.g., fleas, mites and wasps (Okwute 2012). D-Limonene might be responsible for the high toxicity activity recorded from oils from *C. sinensis* and *A. conyzoides* against the larvae stage of *A. gambiae.* Meanwhile, Precocene I and Caryophyllene were identified as key chemical constituents in oil from *O. gratissimum.* Similarly, Caryophyllene was identified as a major chemical constituent in oils from *O. gratissimum* (Dambolena et al. 2010; Saliu et al. 2011; Joshi 2017). The presence of Precocene I and Caryophyllene in *O. gratissimum* may be responsible for its adulticidal activity on female *A. gambiae*. Furthermore, P-Xylene was identified as a major chemical constituent in oil from *N. tabacum*. This is similar to a report from Kidah (2018) where P-Xylene was identified as the key chemical constituent in oils from *N. tabacum*. This might be responsible for the excellent adulticidal activity of *N. tabacum* on female *A. gambiae.* Based on the interesting reports from this study, there is a need to isolate the key chemical constituents of these oils and evaluate their insecticidal potential on *Anopheles* mosquitoes as well as investigate their mechanism of action.

## Conclusions

Plant oils should be seen as supplements to the current vector control intervention in play in developing countries in other to achieve proposed vector control targets. This study was able to establish the bioactivity and efficacy of the evaluated plants exhibiting, varying knockdown, larvicidal and adulticidal potentials. This can be researched further which can be maximized for vector control purposes.

## Acknowledgements

Gratitude goes to the Chief S.L. Edu Research Grant through the Nigerian Conservation Foundation to be able to carry out a part of this work and the Department of Pharmacology Therapeutics, University of Ibadan for space to carry out plant extractions.

## Author Contribution

Olukayode James Adelaja: Conceptualization, Data Curation, Formal Analysis, Funding Acquisition, Investigation, Visualization, Writing – Original Draft Preparation.

Adedayo Olatubosun Oduola: Conceptualization, Supervision, Resources, Writing – Review & Editing.

Adeolu Taiwo Ande: Supervision, Writing – Review & Editing.

Oyindamola Olajumoke Abiodun: Methodology, Resources, Writing – Review & Editing.

Abisayo Ruth Adelaja: Methodology, Formal Analysis, Writing – Review & Editing.

## Data Availability

All data generated or analysed during this study are available from the corresponding author on reasonable request.

## Declarations

### Ethical Approval

Not applicable for research with invertebrates.

### Consent to participate

Not applicable

### Consent for publication

Not applicable

## References

World Health Organization. Vector Borne Diseases. World Health Organization, Geneva. 2019b; 25

World Health Organization. World malaria report. Global malaria programme World Health Organization, Geneva. 2019a; 1–284.

World Health Organization. World malaria report. Global malaria programme World Health Organization, Geneva. 2021; 1–254.

Oduola AO, Adelaja OJ, Ayiegbusi ZO, Tola M, Obembe A, Ande AT, et al. Dynamics of Anopheline vector species composition and reported malaria cases during the rain and dry season in two selected communities in Kwara state. Nig J Paras. 2016;37(2): 158–164.

Okorie PN, McKenzie FE, Ademowo OG, Bockarie M. Nigeria *Anopheles* vector database: an overview of 100 years’ research. PLoS One. 2011;6(12): e28347.

Obi OA, Nock IH, Adebote DA, Nwosu LC. Ecological and Molecular observations on *Anopheles* species (Diptera: Culicidae) breeding in rock pools on iselbergs within Kaduna State, Nigeria. Mol Ent. 2017; 8(01):1–10.

Omotayo AI, Ande AT, Oduola AO, Adelaja OJ, Adesalu O, Jimoh TR, et al. Multiple insecticide resistance mechanisms in urban population of *Anopheles coluzzii* (Diptera: Culicidae) from Lagos, South-West Nigeria. Acta Tropica. 2022;227: 10629. ISSN 0001-706X

Olayemi IK, Ande AT, Chita S, Ibemesi G, Ayanwale VA, Odeyemi OM. Insecticide susceptibility profile of the principal malaria vector *Anopheles gambiae* s.l. (Diptera: Culicidae) in North-central Nigeria. J Vect bor Dis. 2011;48(2): 109–112.

Youmsi RDF, Fokou PVT, Menkem EZ, Bakarnga-Via I, Keumoe R, Nana V, Boyom FF. Ethnobotanical survey of medicinal plants used as insects’ repellents in six malaria endemic localities of Cameroon. J Ethnobio & Ethnomed. 2017; 13(1): 33.

Ande AT, Adelaja OJ, Agada AV, Akanni OI, Omotayo AI. Biological Fitness Costs Associated with the Various Permethrin Resistance Development Statuses in *Anopheles gambiae* in Nigeria. Sri Lanka J Bio. 2022; 7(1): 1–11. http://doi.org/10.4038/sljb.v7i1.73

Oduola AO, Abba E, Adelaja OJ, Ande AT, Yoriyo KP, Awolola TS. Widespread Report of Multiple Resistance in *Anopheles gambiae* Mosquitoes in Eight Communities in Southern Gombe, North East Nigeria. J Arth Bor Dis. 2019.

Adelaja OJ, Oduola AO, Abiodun OO, Adeneye AK, Obembe A. Plants with insecticidal potential used by ethnic groups in North-Central Nigeria for the management of hematophagous insects. Asian J Ethnobio. 2021;4: 65–75.

Ogban EI, Adelaja OJ, Ozovehe AS. Repellent potentials of *Securidaca longepedunculata* Fresen (Polygalaceae) crude extract and essential oil against mosquitoes. Int J Mosq Res. 2020; 7(5): 07–11.

Karunamoorthi K, Ilango K, Endale A. Ethnobotanical survey of knowledge and usage custom of traditional insect/mosquito repellent plants among the Ethiopian Oromo ethnic group. J Ethnopharm. 2009; 125: 224–229.

Kamaraj C, Rahuman AA, Bagavan A, Abduz ZA, Elango G, Kandan P, et al. Larvicidal efficacy of medicinal plant extract against *Anopheles stephensi* and *Culex quinquefasciatus* (Diptera: Culicidae). Trop Biomed. 2010; 27: 211–219.

Ileke KD, Ogungbite OC. *Alstonia boonei* De Wild oil extract in the management of mosquito (*Anopheles gambiae)*, a vector of malaria disease. J Coast Lif Med. 2015; 3(7): 557–563.

Ivoke N, Okafor FC, Owoicho LO. Evaluation of Ovicidal and Larvicidal effects of leaf extracts of *Hyptis suaveolens* (L) Poit (Lamiaceae) Against *Anopheles gambiae* (Diptera: Anophelidae) Complex. Ani Res Int. 2009; 6(3): 1072–1076.

Mgbemena IC. Comparative Evaluation of Larvicidal Potentials of Three Plant Extracts on *Aedes aegypti*. J Amer Sci. 2010; 6(10): 435–440.

Okigbo RN, Okeke JJ, Madu NC. Larvicidal effects of *Azadirachta indica*, *Ocimum gratissimum* and *Hyptis suaveolens* against mosquito larvae. J Agric Tech. 2010; 6(4): 703–719.

Musa AO, Sow GJ, Nasir IF, Shuaibu BU, Edogbanya PRO. Effects of aqueous and methanolic leaf extracts of *Ageratum Conyzoides* l. and *Guiera senegalensis* L. against mosquito larvae in Zaria. J Trop Biosci. 2015; 10: 38–44.

Nzelibe HC, Chintem DGW. Larvicidal Potential of Leaf Extracts and Purified Fraction *Ocimum gratissimum* against *Culex quinquefasciatus* Mosquito Larva. Int J Sci Res. 2015; 4(2): 2254–2258.

Owoeye JO, Ibitoye OA. Analysis of Akure urban land use change detection from Remote Imagery Perspective. USR. 2016; 16: 1–9.

Ayange-kaa AB, Hemen TJ, Onyezili N. The effect of dried leaves extract of *Hyptis suaveolens* on various stages of mosquito development in Benue State, Nigeria. IOSR-JPBS. 2015; 10(6 II): 28–32.

Gillies MT, De Meillon, B. The Anophelinae of Africa South of the Sahara (Ethiopian Zoogeographical Region). Publication of South Africa Institute of Medical Research. 1968; 54: 203–207.

Abbott WS. A method of computing the effectiveness of an insecticide. J Econ Entom. 1925; 18: 265–267.

World Health Organization. Guidelines for laboratory and field testing of mosquito larvicides. 2005; 15–102.

World Health Organization. Susceptibility Test Manual Instructions for Determining the Susceptibility or Resistance of Mosquito Larvae to Insecticides. 2013; WHO/VBC/75.583.

Dua VK, Alam MF, Pandey AC, Rai S, Chopra AK, Kaul VK, et al. Insecticidal activity of *Valeriana jatamansi* (Verbenaceae) against mosquitoes. J Am Mosq Control Assoc. 2008; 24: 315–8.

Finney DJ. “Probit Analysis,” Cambridge University Press, Cambridge, 1971.

Ghosh N, Chowdhury G. Plant extract as potential mosquito larvicide. Ind J Med Res. 2012; 135(5): 581–598.

Bobbo AA, Pukuma MS, Qadeer MA. Assessment of Larvicidal Activity of *Hyptis suaveolens* and *Balanites aegyptiaca* Leaves and Root Extracts against Mosquito Species. IJSRP. 2016; 6(3):10–14

Kavendan K, Mahesh PK, Subramaniam J, Murugan K, William JS. Larvicidal activity of indigenous plant extracts on the rural malarial vector, *Anopheles culicifacies* Giles. J Entom Acar Res. 2014; 46:1747.

Awolola TS, Okwa OO, Hunt RH, Ogunrinade AF. Dynamics of the malaria vector population in coastal Lagos, south-western, Nigeria. Ann Trop Med Parasi. 2002; 96(1): 75–82.

Ahmad MM, Rehman S, Iqbal Z, Anjum FM, Sultan JI. Genetic variability to essential oil composition in four *Citrus* fruit species. Paki J Bota. 2006; 38: 319–324.

Espina L, Somolinos M, Lorán S, Conchello P, García D, Pagán R. Chemical composition of commercial Citrus fruit essential oils and evaluation of their antimicrobial activity acting alone or in combined processes. Food Cont. 2011; 22: 896–902.

Okwute SK. Plants as Potential Sources of Pesticidal Agents: A Review. Pest Advan Chem Botan Pest. 2012; 207–232.

Usman LA, Zubair MF, Olawore NO, Mohammed NO, M’Civer FA, Ismaeel RO. Chemical constituents of flower essential oil of *Ageratum conyzoides* growing in Nigeria. Elixir Org Chem. 2013; 54: 12463–12465.

Karr LL, Coats JR. Insecticidal properties of d-limonene. J Pest Sci. 1988; 13: 287–290

Lee S, Peterson CJ, Coats JR. Fumigation toxicity of monoterpenoids to several stored product insects. J Stored Prod Res. 2003; 39: 77–85.

Almeida MLS, Oliveira AS, Rodrigues AA, Carvalho GS, Silva LB, Lago JHG, et al. Antitermitic activity of plant essential oils and their major constituents against termite *Heterotermes sulcatus* (Isoptera: Rhinotermitidae). J Med Plants Res. 2015; 9: 97–103.

Dambolena JS, Zunino MP, Lopez AG, Rubinstein HR, Zygadlo JA. Essential oils composition of *Ocimum basilicum* L. and *Ocimum gratissimum* L. from Kenya and their inhibitory effects on growth and fumonisin production by *Fusarium verticillioides*. Innova Food Sci Emerg Tech. 2010; 11: 410–414.

Saliu BK, Usman LA, Sani A, Muhammad NO, Akolade JO. Chemical composition and anti bacterial activity of leaf essential oil of *Ocimum gratissimum* grown in North-central Nigeria. Inter J Curr Res. 2011; 33: 022–028

Joshi RK. GC – MS Analysis of the Essential Oil of *Ocimum gratissimum* L. Growing Desolately in South India. Acta Chromatographica. 2017; 29: 111–119.

Kidah MI. Chemical Constituents of the Essential Oil Extracted from *Nicotiana tabacum* Leaves. Biotech J Int. 2018; 21(1): 1–4.

